# Resolving intrinsic ambiguities of the fMRI signal in the human brain

**DOI:** 10.1101/833681

**Authors:** Eulanca Y. Liu, Frank Haist, David Shin, Richard B. Buxton

## Abstract

Functional magnetic resonance imaging (fMRI) using the blood oxygenation level dependent (BOLD) signal is a standard tool in human neuroscience studies. However, the BOLD signal is physiologically complex, depending on the baseline state of the brain and the balance of changes in blood flow and oxygen metabolism in response to a change in neural activity. Interpretation of the magnitude of the BOLD response is thus problematic; specifically, group differences of the BOLD response to a standard task or stimulus, or a change after administration of a drug, could be due to a change in the neural response, neurovascular coupling, or the baseline state. While changes in oxygen metabolism can serve as a biomarker of neural activity change, reflecting the energy cost of activity, oxygen metabolism cannot be estimated from BOLD measurements alone. Here we used the effects of caffeine as a test case to show that a suite of additional noninvasive measurements, requiring no manipulation of inhaled gases, makes it possible to untangle the underlying effects of caffeine on blood flow and oxygen metabolism. After caffeine administration, the BOLD response to a standard motor task was reduced, but oxygen metabolism was not reduced; rather, caffeine reduced the balance of blood flow relative to oxygen metabolism, both in the baseline state and in response to the task. The MRI methods used required only an additional 12 minutes of scanning, can be implemented on any MRI system, and painted a more complete picture of the physiological effects of caffeine. In addition to the assessment of drug effects in the human brain, these methods make it possible to resolve ambiguities of the BOLD signal in studies of development, aging, and disease. (Duplicated from first paragraph of manuscript)

---

Functional magnetic resonance imaging (fMRI) using the blood oxygenation level dependent (BOLD) signal is a standard tool in human neuroscience studies. However, the BOLD signal is physiologically complex, depending on the baseline state of the brain and the balance of changes in blood flow and oxygen metabolism in response to a change in neural activity. Interpretation of the magnitude of the BOLD response is thus problematic; specifically, group differences of the BOLD response to a standard task or stimulus, or a change after administration of a drug, could be due to a change in the neural response, neurovascular coupling, or the baseline state. While changes in oxygen metabolism can serve as a biomarker of neural activity change, reflecting the energy cost of activity, oxygen metabolism cannot be estimated from BOLD measurements alone. Here we used the effects of caffeine as a test case to show that a suite of additional noninvasive measurements, requiring no manipulation of inhaled gases, makes it possible to untangle the underlying effects of caffeine on blood flow and oxygen metabolism. After caffeine administration, the BOLD response to a standard motor task was reduced, but oxygen metabolism was not reduced; rather, caffeine reduced the balance of blood flow relative to oxygen metabolism, both in the baseline state and in response to the task. The MRI methods used required only an additional 12 minutes of scanning, can be implemented on any MRI system, and painted a more complete picture of the physiological effects of caffeine. In addition to the assessment of drug effects in the human brain, these methods make it possible to resolve ambiguities of the BOLD signal in studies of development, aging, and disease.

The cerebral metabolic rate of oxygen (CMRO_2_) reflects energy generation from oxidative metabolism of glucose. It is dominantly driven by processes related to neural signaling, primarily the cost of restoring sodium and calcium gradients across cellular membranes following excitatory synaptic activity^2^. Oxygen is delivered to the tissue by cerebral blood flow (CBF) with the balance of CMRO_2_ and CBF rates reflected by the oxygen extraction fraction (OEF), or the fraction of arterial blood oxygen extracted in passing through the capillary bed and metabolized in the tissue mitochondria. Empirically, the OEF is relatively uniform in the resting human brain with a value of about 0.4 ^3^. The essential yet unexpected physiological phenomenon is that when neural activity increases in response to a task or stimulus, the fractional CBF change is much larger than the fractional CMRO_2_ change, by a factor of two or more, with the seemingly paradoxical effect of reducing the OEF when CMRO_2_ increases^4^. The biological function served by the large CBF increase may be to prevent a fall in the tissue oxygen concentration, although current evidence indicates that dynamic CBF modulation is controlled in a feedforward manner by mechanisms related to neural activity, rather than direct feedback from oxygen levels^5-7^. Taking CMRO_2_ as a biomarker of the energy cost of neural activity, the coupling ratio of CMRO_2_ and CBF changes can be taken as a biomarker of neurovascular coupling.

The BOLD signal change following a change in neural activity is not a direct indicator of a change in CMRO_2_, but instead reflects the change in the deoxyhemoglobin content of tissue, with the mismatch of fractional changes in CMRO_2_ and CBF modulating the baseline deoxyhemoglobin level^8^. Due to its paramagnetic effects, deoxyhemoglobin reduces the MR signal, and during neural activation the reduced OEF lowers the deoxyhemoglobin content and increases the MR signal. The magnitude of the BOLD response to a change in neural activity thus depends on the CMRO_2_ change, the degree of neurovascular coupling that determines the CBF change, and baseline deoxyhemoglobin content of tissue. The goal of quantitative fMRI is to use additional measurements to untangle this physiological complexity and ideally produce measurements of absolute CBF and CMRO_2_ in both the baseline and activation states^9,10^. A primary tool is arterial spin labeling (ASL), which makes it possible to measure dynamic CBF and BOLD responses simultaneously^11,12^ (see Methods). For quantitative CMRO_2_ measurements, though, measurements of baseline OEF and deoxyhemoglobin content also are needed. A current approach is to measure the CBF and BOLD responses to altered gases, hypercapnia and hyperoxia, requiring a special experimental setup for gas delivery^13-15^. Because of the additional complexity of this experiment, the gas method is used only in dedicated research labs. In addition, the gas methods bring additional challenges in accurately modeling the physiological responses.

In the current study we complemented the ASL/BOLD experiment with two different MRI methods that do not require the subject to breathe special gases, but are sensitive to the additional needed physiological parameters of the baseline state^16^. As with the BOLD effect, the physical basis of these tools is the effect of deoxyhemoglobin on the relaxation rate of the precessing transverse magnetization that generates the MR signal. For the BOLD signal, the appropriate relaxation rate is tissue R_2_*, which is reduced when OEF is reduced^8^. The quantitative methods used here exploit two additional relaxation rates: R_2_′ of tissue in the baseline state—essentially the difference between R_2_* and the relaxation rate R_2_ measured with a spin echo experiment—which is sensitive to baseline deoxyhemoglobin content^17-20^; and R_2_ of venous blood, which is sensitive to baseline OEF^21,22^ (see Methods). This suite of quantitative tools was used to measure baseline CBF and CMRO_2_ and the activation changes in CBF and CMRO_2_ to a simple finger-tapping task before and after administration of 200mg of caffeine in healthy human subjects. Our goal was to use caffeine as a test case in which the physiological effects are somewhat known from previous studies, and address the central question: can these noninvasive quantitative methods untangle both baseline and activation effects that otherwise make the BOLD response to a stimulus ambiguous?

The administration of caffeine in the baseline non-task condition had significantly different effects on CBF and CMRO_2_ within superior cortical grey matter (Figure 1A), with caffeine strongly reducing mean CBF (21.9% reduction) with no significant change in mean CMRO_2_ (7.0% reduction). Previous studies have consistently found similar levels of reduced CBF, but results for CMRO_2_ have varied^11,23-27^. The CBF decrease (absolute change in the value) with caffeine was consistent across subjects regardless of their baseline CBF (Figure 1B). We also found increased variance in OEF across the subjects following caffeine ingestion (Figure 1C), leading to a larger variance in CMRO_2_ (Figure 1D) calculated from the OEF and CBF measurements; this suggests that CMRO_2_ effects may be more complicated in a way that contributes to variability of CMRO_2_ results observed across different studies. The parameter R_2_′ was relatively unchanged, possibly due to a partial cancelling of effects of increased OEF, tending to increase total deoxyhemoglobin, and reduced flow, tending to decrease total deoxyhemoglobin.

**Figure 1:**
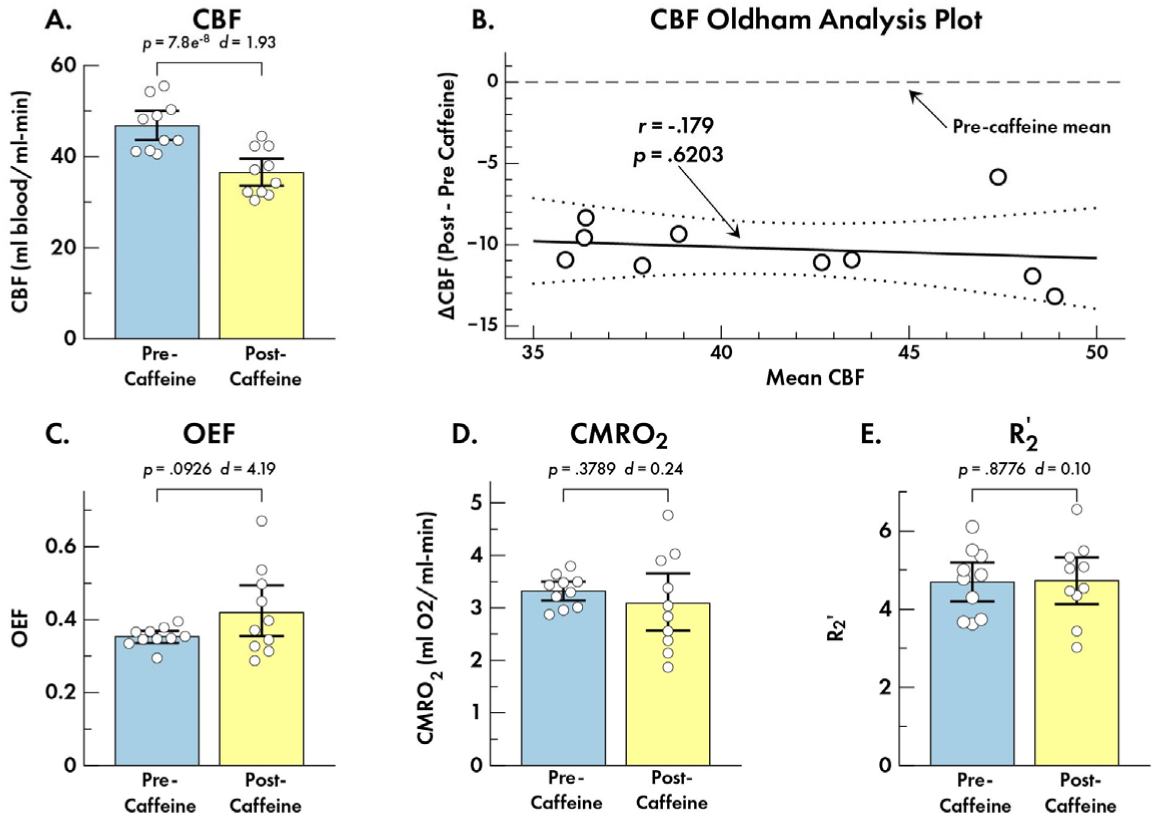
Baseline pre- and post-caffeine physiological variables averaged over a single slice in superior cortical grey matter. (**A**) Blood flow (CBF) decreased after caffeine administration in all subjects (caffeine condition x physiology interaction: *p* = 2.2e^-7^, Cohen’s *d* = 9.21). (**B**) The lack of a correlation between the difference and the average of the pre- and post-caffeine values, applying Oldham’s analysis^1^, is consistent with a uniform absolute shift of CBF across subjects independent of the pre-caffeine CBF value. (**C**) The OEF increased slightly and showed a large increase in variance across subjects. (**D**) Oxygen metabolism (CMRO_2_) was similar pre- and post-caffeine, although the variance across subjects was sharply increased, driven by the large increase in variance of OEF, which is used to calculate baseline CMRO_2_. (**E**) The parameter R_2_′ showed little change post-caffeine. Error bars = 95% confidence intervals.

We evaluated the effect of task activation across physiological measures within a task-sensitive region of interest (ROI) in motor cortex in pre-caffeine and after caffeine states (Figure 2). As in the larger grey matter region, CBF and CMRO_2_ responded differently within this ROI at baseline (Figure 2A), with CBF declining after caffeine (17.5% reduction) but no significant change observed in CMRO_2_ (4.9% reduction). These two physiological measures also responded differently during task performance (Figure 2B) with absolute mean CBF change during task performance reduced by 29.0% after caffeine administration and absolute CMRO_2_ change increased by 23.4%, although this latter difference was not significantly different between the conditions. The activation-related BOLD response decreased after caffeine administration by 27.7% (Figure 2C). Interestingly, the neurovascular coupling ratio, taken as the ratio of the fractional CMRO_2_ change to the fractional CBF change in response to the task (Figure 2C), showed a robust 47.7% increase.

**Figure 2.**
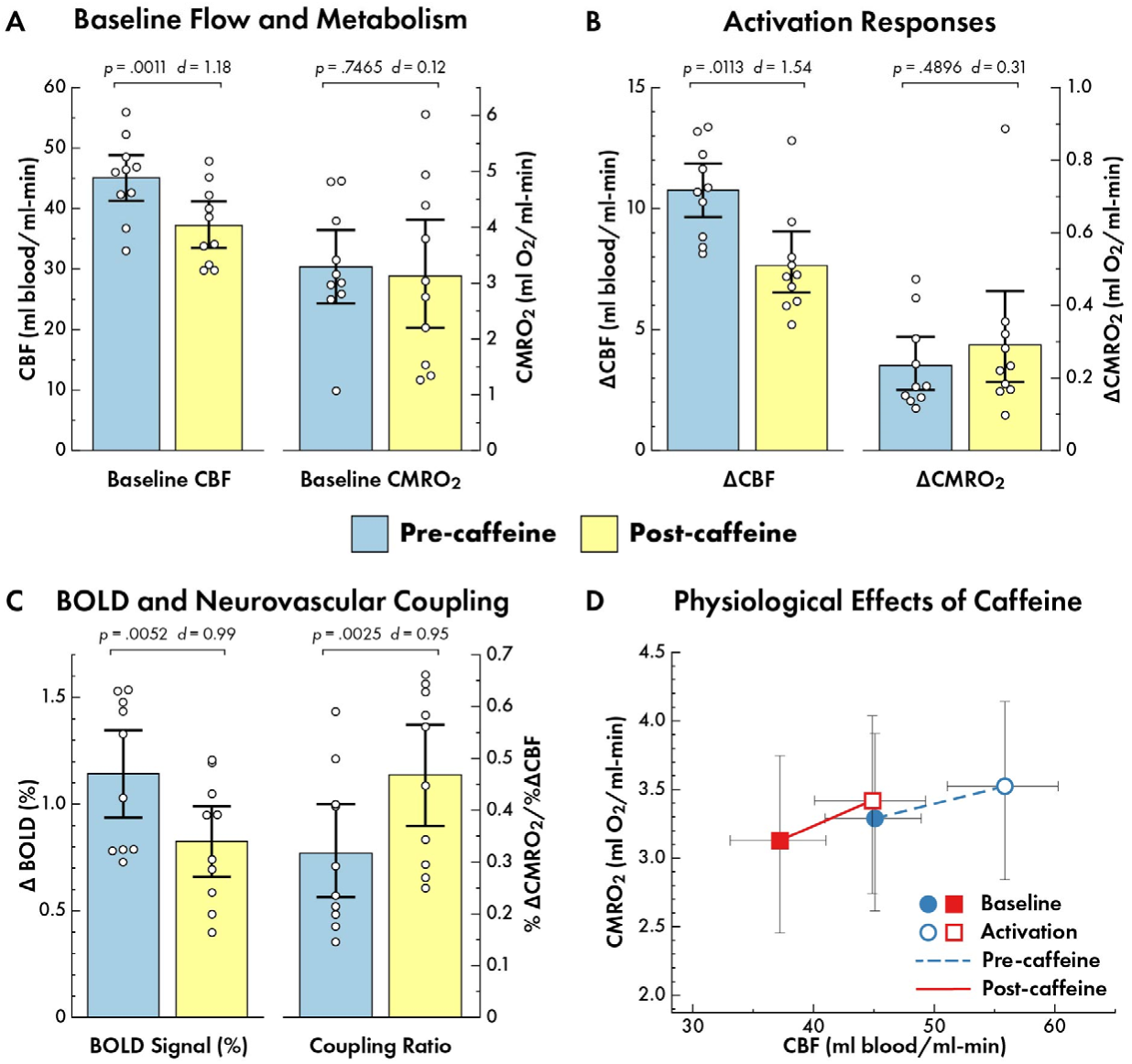
Pre- and post-caffeine baseline and activation physiological variables measured in primary motor cortex ROI. Results are for a small ROI (volume 0.74+/-0.39 cm^3^) activated by the motor task. (**A**) The effect of caffeine on the baseline state was a reduction of CBF with no change in CMRO_2_. The CBF and CMRO_2_ responded significantly differently (caffeine condition x physiology: F_1,9_ = 26.63, *p* = .0006, Cohen’s *d* = 3.44). (**B**) The effect on activation responses to the motor task was a reduction of the absolute CBF response and an increase of the CMRO_2_ response, although the latter was not statistically significant. The CBF and CMRO_2_ responded significantly differently (caffeine condition x physiology: F_1,9_ = 25.18, *p* = .007, Cohen’s *d* = 2.32). (**C**) However, the neurovascular coupling ratio—the ratio of the fractional change in CMRO_2_ to the fractional change in CBF—was strongly increased following caffeine, while the BOLD response to the motor task was strongly reduced. Note that the reduction of the BOLD signal was driven by this change in neurovascular coupling, not by a reduction of the CMRO_2_ response to the task. (caffeine condition x physiology: F_1,9_ = 20.55, *p* = .0014, Cohen’s *d* = 3.02). (**D**) Comparing the absolute values for CBF and CMRO_2_ in each of the states (pre-/post-caffeine, baseline/activation) suggests that activation and caffeine effects move along a common activation/deactivation path in the CBF/CMRO_2_ plane. Error bars = 95% confidence intervals.

This study demonstrates the feasibility of using additional MRI measurements to untangle the physiological baseline and activation effects underlying current fMRI signals. Here we found a reduced BOLD response to a standard task after administration of caffeine, but this result alone could not be interpreted in a meaningful way in terms of a change in brain physiology—even the direction of the BOLD change alone was misleading in relation to the drug effect on the CMRO_2_ change to the stimulus. The additional measurements revealed that the primary effect of caffeine was to reduce the balance of CBF relative to CMRO_2_, both in the baseline state and in the fractional changes in response to the task, with no evidence for a reduction in the energy cost of performing the task. That is, BOLD signal change to the task after caffeine was largely driven by the effects of a strong change in neurovascular coupling (Figure 2C), rather than a change in the neural response.

Finally, these data illustrate that full quantitative measurements of CBF and CMRO_2_ in both the baseline and activation states (Figure 2D) make it possible to examine the dynamics across different states in a way that is not possible with BOLD imaging alone, which only measures a change between two conditions. The interesting finding in comparing absolute CBF and CMRO_2_ values across caffeine and neural activation manipulations is that these physiological variables are similar in the pre-caffeine baseline state and the post-caffeine activation state (blue solid circle vs. red open square in Figure 2D), suggesting the possibility that the separate effects of caffeine and neural stimulus may move CBF and CMRO_2_ along a common curve in the flow-metabolism plane.

The finding of distinct effects of caffeine on CBF and CMRO_2_ is consistent with caffeine acting as an antagonist at adenosine receptors^28^. Adenosine has both vascular and neural effects, acting as a strong vasodilator while tending to inhibit neural activity. Blocking adenosine receptors should then have a strong effect of reducing CBF, with perhaps an accompanying effect of increasing CMRO_2_ if there is a net reduction in inhibition. However, the latter effect is harder to predict given the possible roles of several types of adenosine receptor. Though further work is needed to understand the full physiological effects of caffeine on CBF and CMRO_2_, these results support the ability to probe these mechanisms with quantitative fMRI tools.

The methods used here are representative of approaches being developed to exploit the effect of deoxyhemoglobin on different relaxation rates of the MR signal^21,29,30^, and our current results demonstrate that with these methods, the ambiguities of the BOLD response can be untangled in a relatively small cohort of subjects and with only a modest increase in imaging time over a standard BOLD-fMRI study. Continued development of these tools is likely to lead to improvements in sensitivity and the possibility of clinical applications^29^ (the potential for individual measurements of brain state is evaluated in more detail based on the current data in Supplementary Information). The current study demonstrates the capability of these methods to paint a much more detailed and quantitative picture of brain function than is possible with BOLD measurements alone, particularly for interpreting BOLD response differences between groups in studies of development and disease or after an intervention.

## Methods

### 1. Subjects

Ten healthy adults (5 female, mean age=23.5 years, range 21-28) participated in this study. All subjects reported drinking the equivalent of 12 ounces or more of black coffee per day. We asked subjects to abstain from ingesting caffeine for at least 4 hours prior to the start of the imaging session. The study was approved by the University of California, San Diego Human Research Protections Program. Written informed consent was obtained from all subjects; all were remunerated for their participation.

### 2. Overview of the experimental design

The 3-hour MR scan session included: 1) a suite of MRI measurements during a “pre-caffeine” condition; 2) administration of a 200-mg caffeine tablet followed by a 30 min break; and 3) the same suite of MRI measurements repeated during a “post-caffeine” condition. The pre- and post-caffeine conditions included one scan to measure tissue R_2_′ and one to measure venous blood R_2_, both collected during a “fixation only” resting condition, and either two (pre-caffeine) or one (post-caffeine) functional runs with simultaneous measurement of CBF and BOLD signal changes, the latter measured as a change in tissue R_2_*. Subjects performed a sequential finger-tapping task cued via a visual stimulus administered in a blocked-design format during the functional run. We will refer to the fixation-only rest periods between finger-tapping blocks as baseline. Physiological changes during finger-tapping activation states were calculated relative to this baseline.

We analyzed data from a single axial slice that included the primary motor area in two ways. First, we tested the consistency of the noninvasive baseline measurements with predictions from previous studies of caffeine effects within the mean grey matter (GM) data from the entire slice (Figure 1). Next, we tested for combined baseline and activation effects to paint a full picture of absolute CBF and CMRO_2_ in the baseline and activation states within a region of interest (ROI) in primary motor cortex (Figure 2). The ASL data provided quantitative CBF values for both states. Baseline venous blood R_2_ was used to estimate the oxygen extraction fraction (OEF). Baseline CMRO_2_ was estimated as the product of OEF, baseline CBF, and an assumed value of arterial O_2_ concentration of 9 mM. The fractional change in CMRO_2_ to the task was estimated as in a calibrated BOLD experiment by using the measured dynamic CBF/BOLD changes to the task and taking R_2_′ * TE as the estimate of the scaling factor *M* in the Davis model of the BOLD signal.

### 3. Functional task

Visual cues for the task were projected onto a screen viewed by the subject through a mirror attached to the head coil. The motor-visual task used a blocked-design format that included alternative 33.6 sec periods of visual-motor task blocks and fixation-only rest blocks. Thus, each run collected three blocks each of task and rest. Motor-visual task blocks used a patterned finger-tapping task paced by a red circle that was presented at the center of the field of view at 2 Hz. During the task, subjects pressed one of four buttons on a 4-button box with their right hand in the pattern 1-3-2-4 or index-ring-middle-pinky. Surrounding the red circle was a radial flickering checkerboard. Subjects were instructed to ignore the flickering checkerboard and focus on the red circle to cue their button-presses in the correct pattern. We also analyzed responses from the visual cortex (see Supplementary Information). The responses to the unattended visual stimulus were less robust than the finger-tapping responses. During the resting blocks, subjects were instructed to keep eyes open and fixate on a black cross presented at the center of the field of view. The cross turned from black to red 2 sec before the beginning of the motor block to signal the transition to the task. In addition, we collected 2 min and 30 sec of fixation-only rest prior to the beginning of the six task-rest blocks. Two functional runs were acquired in the pre-caffeine state, the second run immediately following the first run, and a single functional run was acquired in the post-caffeine state. The repetition time (TR) of the image acquisition was 4.2s.

### 4. Functional imaging: BOLD and CBF dynamics

#### 4.1. Image acquisition

We scanned using a General Electric (GE) Discovery MR750 3.0T MRI system with a three echo pseudo-continuous arterial spin labeling (pCASL) pulse sequence and an eight-channel receive-only head coil^31,32^. Frames within a task run alternated between label and control images. We calculated the dynamic CBF time series from the difference of tag and control images for the shortest echo time (TE) images, and a BOLD time series from the tag and control average. The PCASL sequence (TR = 4.2s) consisted of 1650ms labeling duration, followed by 1600ms post labeling delay during which two background suppression pulses were applied (1580ms and 472ms relative to the first excitation pulse) to achieve 50% tissue suppression at the middle of the slice pack^31,32^. We obtained 12 axial slices (5 mm thick/1 mm gap, FOV=256mm, matrix 64×64) using single-shot spiral readouts. Three echoes were collected per slice (TE=3.3ms/25ms/46.7ms). Two calibration and reference scans were acquired with the same slice prescription as that of the BOLD-ASL sequence in both the pre-and post-caffeine states: a field map to correct distortions in the spiral acquisition due to magnetic field inhomogeneity^33^ (eight-shot spiral acquisition, TE=11ms, TR=2000ms). We also collected a cerebral spinal fluid (CSF) reference scan for CBF quantification that used a single-echo spiral EPI acquisition (TE=3.3ms, TR=4000ms)^11,34^, and a minimum contrast (min/con) scan that we used to correct for transmit and receiver coil inhomogeneities^35^ in the ASL images (eight-shot spiral acquisition, TE=11ms, TR=2000ms). We acquired a high-resolution T1-weighted anatomical scan (FSPGR) for image registration and tissue segmentation in both the pre- and post-caffeine conditions for each session.

#### 4.2. Image preprocessing

Data processing was done as described by Liu et al^16^. Field map correction was first performed on raw ASL data to account for magnetic field inhomogeneities^33^. The functional scans were motion corrected and registered to one image in each of the runs using AFNI software^36^. The first four images were dropped to allow the signal to have reached a steady state. The ASL data were then corrected for coil sensitivity inhomogeneity using the min/con images^35^. Surround-subtraction was applied to first-echo ASL images to produce CBF-weighted images, then converted to physiological CBF units using the CSF scan as reference^34^. Surround-averaging was applied to the first- and second-echo images; these values were used to calculate BOLD R_2_*-weighted images. For each time series, after ROI selection (described below), the mean R_2_* was subtracted to form ΔR_2_*(t) time series, which were then converted back to δBOLD signal without systematic effects of drift^16^.

### 5. Baseline imaging: OEF and CMRO_2_

#### 5.1 VSEAN acquisition

In each of the two baseline states, a measurement of venous oxygenation was made using Velocity-Selective Excitation and Arterial Nulling (VSEAN)^22^ developed by Guo and Wong and tested in a previous study^16^. The VSEAN pulse sequence is applied on one slice at a time, and works by combining arterial nulling and a velocity selective module to isolate the signal of venous blood, and then measuring the signal decay rate R_2_ with a spin-echo module using three effective echo times of 25, 50 and 75 ms. Other pulse sequence parameters were: FOV = 256 mm × 256 mm, single-slice spin echo with spatial-spectral excitation, two slice-selective hyperbolic secant refocusing pulses, single-shot spiral readout, v_e_=2 cm/s in slice-selective direction, slice-selective post-saturation pulses, 81 acquisitions preceded by two dummy scans and including six “cos” modulated reference scans^22^. The total scan time for a single slice acquisition was 4 min.

#### 5.2 Calculation of baseline OEF

The venous blood R_2_ values were translated to blood oxygen saturation of hemoglobin Y_v_ via a R_2_-Y_v_ calibration curve based on an analysis of bovine blood samples measured *in vitro* at 3 Tesla^37^. The equation for the curve used in this analysis is^22^ 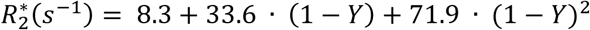. The OEF was then taken as 1-Y_v_, based on the assumptions that arterial hemoglobin was fully saturated with oxygen and dissolved O_2_ is negligible compared to hemoglobin-bound O_2_. Detailed acquisition and analysis parameters were as reported in a previous manuscript^16^.

#### 5.3 Calculation of baseline CMRO_2_

The rate of O_2_ delivery per ml of tissue at baseline is the product of CBF and the total arterial oxygen concentration C_a_O_2_, and CMRO_2_ is then the rate of delivery times the OEF. The equation to determine total arterial oxygen concentration is C_a_O_2_=1.36(Hb)(S_a_O_2_/100)+0.0031(P_a_O_2_), with units as follows: C_a_O_2_ in ml/dL, 1.36 ml O_2_/1 g Hb, Hb in g/dL, S_a_O_2_ in %, 0.0031 g/dl/mmHg, and P_a_O_2_ in mmHg. Hemoglobin concentration was assumed according to mean healthy ranges set by UC San Diego Health: for females, [Hb]=14g/dL; males, [Hb]=15.7g/dL. The age of each subject was also factored into CaO_2_ determination through the estimate of P_a_O_2_ (Estimated normal P_a_O_2_ ≅ 100 mmHg–(0.3) age in years). CBF is expressed in ml/100ml/min, and C_a_O_2_ from this equation is divided by 100 to give CMRO_2_ units in ml/100ml/min. CMRO_2_ in ml/100ml/min can be converted to mM/min (1 mL O_2_ at 310 K, 1 atm = 0.03933 mmol O_2_) For reference, 1 mM/min=2.54 ml O_2_/100ml/min=2.42 ml O_2_/100g/min = 100 μmol/100ml/min = 95 umol/100g/min, with assumed density of blood = 1.05 g/mL at 310 K.

### 6. Baseline imaging: Tissue R_2_′

#### 6.1 FLAIR-GESSE acquisition

Tissue R_2_′ was measured with a method called FLuid Attenuated Inversion Recovery-Gradient Echo Sampling of Spin Echo (FLAIR-GESSE)^20^. The prescription used for this sequence matched that of the BOLD-ASL acquisition, but with a 2mm thick slab and 4 mm gap. Two pairs of GESSE image series were acquired separately, one as an early spin echo series (63.6 ms after excitation) and one as a late spin echo series (83.6 ms after excitation). This provides data on opposite sides of the spin echo but collected at the same absolute time after the initial excitation, which helps correct for problems with multiple R_2_ decay rates within a voxel^20^. For each series, 64 samples of each decay curve were collected asymmetrically. CSF nulling was applied in a FLAIR module (TR=3.5s, TI=1.16s) to the center slices to minimize CSF contamination of R_2_′ measurements. Field offsets were acquired that were used to calculate large-scale field inhomogeneities and remove their contribution to the R_2_′ estimate^38^. R_2_′ was calculated as described in Simon et al^20^. Total acquisition time was 8 min.

### 7. Data analysis: Grey matter baseline effects of caffeine

#### 7.1 Grey matter (GM) mask

The BOLD and ASL time courses were upsampled from 64×64 to 256×256 matrix size by zero-padding in the Fourier domain with the original data centered in k-space, then reconstructing the image again through an inverse Fourier transform. A GM mask was created using a threshold for CBF of 35 ml/100ml/min for baseline and caffeine-free CBF, and 20 ml/100ml/min for post-caffeine CBF. The lower threshold for the post-caffeine data was chosen to compensate for the expected CBF lowering effect of caffeine and thus produce approximately the same number of mask voxels as the pre-caffeine study. The data and masks were smoothed using a Gaussian smoothing kernel with a full width at half maximum of 8 voxels at the upsampled resolution (2 voxels in terms of the original 64×64 resolution).

#### 7.2 Baseline CBF

The masks were applied to the CBF measures across the entire time series. Subsequent measurements of the various baseline parameters were determined by averaging over the data from the 2.5 min baseline resting period. We calculated baseline CBF from the average CBF value over this resting period for each voxel in the respective region, then took the mean across all voxels in the ROI. We used this process for both pre- and post-caffeine states.

#### 7.3 *Baseline* R_2_′

FLAIR-GESSE data were analyzed per Simon et al^20^. The raw GESSE data were first upsampled to the same 256×256 matrix size as the BOLD/ASL. R_2_′ values were fitted to the raw data for each voxel; we used the median of the GM mask ROI as the estimate of ROI R_2_′.

### 8. Data analysis: Dynamic changes to the task

#### 8.1 Functional ROI selection for motor activation

We defined GM mask ROI voxels using a voxel-wise Pearson correlation between the CBF dynamics and the stimulus timeseries that exceeded a threshold within the left primary motor cortex of *r*≥0.30. Separate functional ROIs were obtained for each of the two states, so that the ROIs were comparable functionally across pre-caffeine and post-caffeine conditions.

#### 8.2. ROI definition caveats

We chose to focus on the approach of functional alignment, rather than anatomical alignment, so all numbers derived from the ROI’s were in the native space of the acquisition in each state. Our concern was that the interpolation involved in anatomical alignment of these relatively coarse images would introduce bias in the calculated physiological responses. That is, registration differences between the ASL and anatomical images could be due to subject motion, but also to image distortions due to the different acquisition techniques that may not be fully corrected with the standard alignment tools. One concern was associated with whether using ROIs selected from the baseline CBF data would lead to a smaller ROI post-caffeine because of the reduced baseline CBF. To avoid this bias, we limited the ROIs to be the same size in both conditions by adjusting the threshold level in the post-caffeine data until the mask had approximately the same number of voxels as the pre-caffeine mask.

#### 8.3 BOLD and CBF task changes

We calculated the task-related fractional change in CBF (δCBF by taking the mean value from the last 21 seconds of each of the 3 task blocks to allow for a task onset transition (12.6 seconds or 3 TRs). Baseline CBF was taken as the mean from the following windows: the 29.4 seconds before the start of the first stimulus, the 29.4 seconds before the start of the second stimulus, the 29.4 seconds before the start of the third stimulus, and 29.4 seconds after the end of the last stimulus, skipping one 4.2 second window (1 TR). That is, one TR was skipped for the window after each of the task blocks ended allowing the transition back to baseline. The fractional task-related CBF change was calculated by taking the difference of the mean baseline and stimulus values and dividing by the baseline value. The BOLD signal change (δBOLD, the fractional change of the MR signal) to the task was calculated by taking the mean value over the same temporal windows. Since the signal is already expressed as a fractional BOLD change after being converted from δR_2_*, the fractional task-related change in BOLD was taken as the difference between the mean values from baseline and activated values.

#### 8.4 Estimation of task-related CMRO_2_

We calculated fractional CMRO_2_ (δCMRO_2_) using the fractional stimulus-related BOLD (δBOLD) and CBF (δCBF) change using the calibrated BOLD method^39^. The fractional BOLD signal is modeled as:

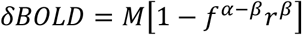

where the physiological changes are *f*=(1+δCBF) and *r*=(1+δCMRO_2_). In the traditional calibrated BOLD approach, values are assumed for the parameters α and β and *M* is estimated by measuring the CBF and BOLD responses to increased CO_2_ in the gas the subject breathes, with the assumption that *r*=1 for mild hypercapnia. In a previous study we found that assuming α=0.2, β=1.3 and *M*= R_2_′TE yielded similar estimates of CMRO_2_ change to a hypercapnia calibration^16^.

### 9. Statistical methods

We used IBM SPSS Statistics v. 26 for all analyses.

#### 9.1 Statistical comparisons of pre- and post-caffeine responses from within the GM ROI

Results from pre- and post-caffeine measures within the GM mask ROI for CBF, OEF, CMRO_2_, and R’_2_ (see Figures 1A, 1C-1E) used paired-sample *t* tests with bootstrap (N=10,000) to calculate 95% confidence intervals (CI) and validate the parametric results. We used Oldham’s approach^1^ to evaluate the relationship between the change in a measurement from pre-to post-caffeine states and the initial (pre-caffeine) value of the measurement. To do this, the mean CBF from the pre- and post-caffeine states are plotted against the change in post-caffeine CBF relative to the pre-caffeine CBF. Oldham’s approach considers the fact that pairs of measures from a subject are correlated and some aspects of the response to a treatment are idiosyncratic and not related to the treatment effect alone and initial high or low values may show regression to the mean. At the group level, Oldham’s approach considers changes in variance across the measures that influence the correlation statistics. We compared mean CBF against the change in CBF using Pearson correlation analysis with bootstrap (N=10,000) to define 95% CI.

#### 9.2 Statistical comparisons of pre- and post-caffeine responses from within the Motor Cortex ROI

Results from pre- and post-caffeine measures within the Motor Cortex mask ROI for CBF, OEF, CMRO_2_, and R’_2_ (see Figures 2A-2C) used repeated measures ANOVA with physiological condition and caffeine state as within-subjects factors with bootstrap (N=10,000) to calculate 95% confidence intervals (CI) and validate the parametric results. Post-hoc analysis used Bonferroni-corrected paired-sample *t* tests with bootstrap (N=10,000) to calculate 95% confidence intervals (CI) and validate the parametric results.

### 10. Data and code availability

The data that support the findings of this study are available from the corresponding author, RBB, upon reasonable request. The data and code sharing adopted by the authors comply with requirements by the National Institutes of Health and by the University of California, San Diego, and comply with institutional ethics approval.

## Acknowledgments

This work was supported by the National Institutes of Health grants NS036722, MH111359, MH112969, and MH113295.

