## Supplementary Information for "Resolving intrinsic ambiguities of the fMRI signal in the human brain"

**Supplementary Information for Liu et al. *Resolving intrinsic ambiguities of the fMRI signal in the human brain***

### S1. Baseline grey matter measurements for two sections containing primary motor area ('Superior GM') and primary visual area ('Caudal GM').

During the motor task the subject was also presented with a flickering checkerboard, but told to ignore the visual stimulus and focus on the motor task. The response in the visual cortex ('Caudal GM') is thus to an unattended stimulus, and the results are less robust than the motor results, while in general supporting the findings in motor cortex ('Superior GM'). Data are shown for the two GM regions together for ease of comparison. The data are summarized in Supplementary Table S1 and shown graphically in Supplementary Figure S1.

**Supplementary Table S1: Average GM changes to caffeine.** Asterisks indicate significant difference from zero with 1-tailed T-testing. *** $p<0.001$; ** $p<0.01$; * $p<0.05$.

| Caudal GM | | | | |
|---|---|---|---|---|
| | CBF (ml/100ml/min) | OEF | $R_2'$ ($s^{-1}$) | $CMRO_2$ (ml/100ml/min) |
| Pre-caffeine | 66.21 | 0.36 | 3.91 | 4.80 |
| Standard deviation | 7.69 | 0.06 | 0.42 | 0.78 |
| CV (%) | 11.62 | 16.14 | 10.64 | 16.36 |
| Post-caffeine | 49.22 | 0.43 | 3.96 | 4.27 |
| Standard deviation | 6.97 | 0.10 | 0.49 | 1.11 |
| CV (%) | 14.15 | 22.73 | 12.27 | 26.04 |
| Δ Post – Pre | -16.99*** | +0.07* | +0.05 | -0.52 |
| Standard deviation | 5.76 | 0.12 | 0.34 | 1.55 |
| % Change from pre-caffeine | -25.53*** | +23.72* | +1.54 | -7.00 |
| **Superior GM** | | | | |
| | CBF (ml/100ml/min) | OEF | $R_2'$ ($s^{-1}$) | $CMRO_2$ (ml/100ml/min) |
| Pre-caffeine | 46.73 | 0.35 | 3.59 | 3.32 |
| Standard deviation | 5.50 | 0.03 | 0.33 | 0.30 |
| CV (%) | 11.76 | 7.70 | 9.18 | 9.13 |
| Post-caffeine | 36.48 | 0.42 | 3.73 | 3.09 |
| Standard deviation | 5.13 | 0.12 | 0.27 | 0.92 |
| CV (%) | 14.07 | 28.63 | 7.25 | 29.90 |
| Δ Post – Pre | -10.25*** | 0.07* | 0.14* | -0.23 |
| Standard deviation | 2.07 | 0.11 | 0.21 | 0.79 |
| % Change from pre-caffeine | -22.04*** | +18.50* | +4.19* | -7.62 |

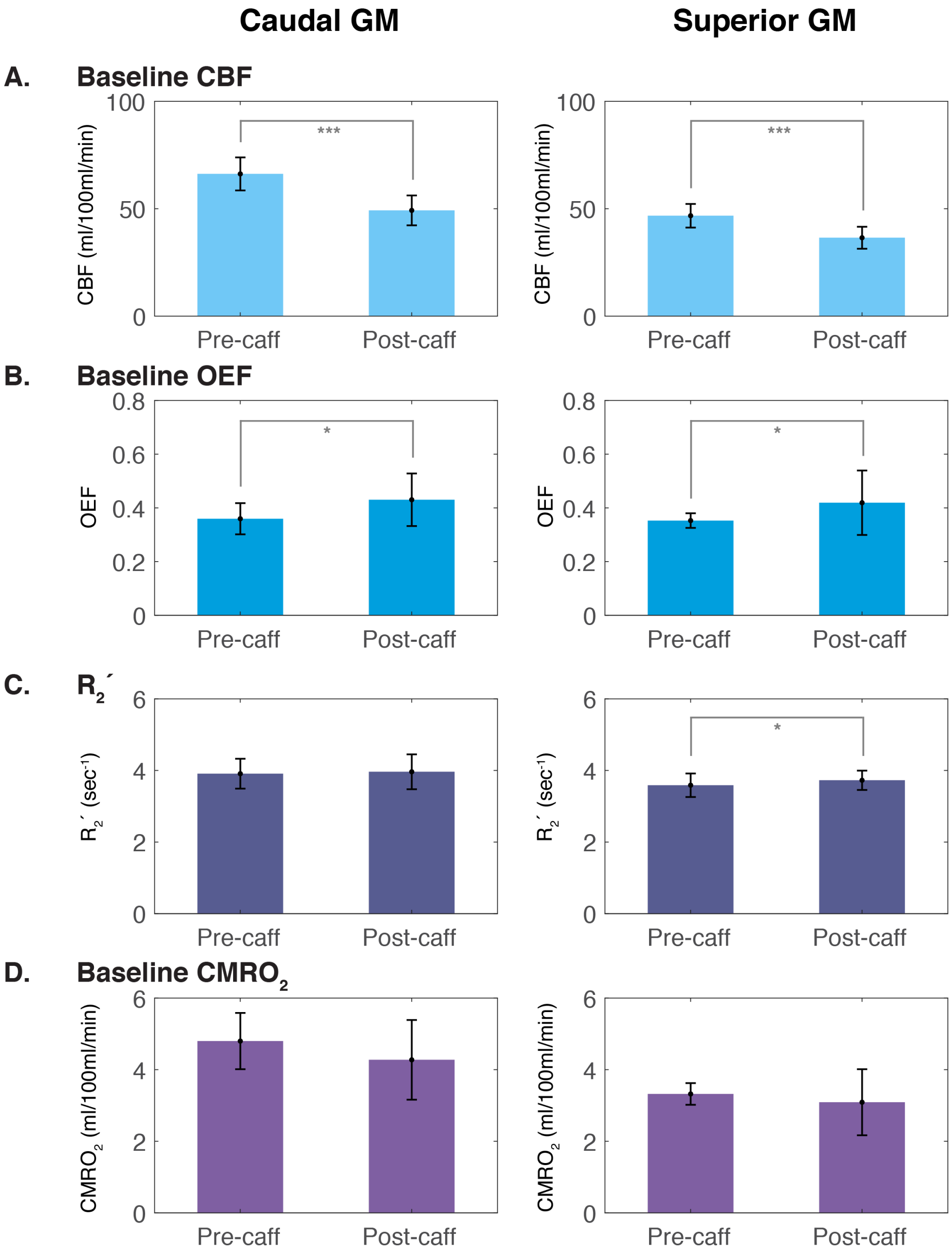

**Supplementary Figure S1: Baseline values pre- and post-caffeine.** Mean values across 10 subjects of (**A**) CBF, (**B**) OEF, (**C**) $R_2'$, and (**D**) $CMRO_2$ in the pre- and post-caffeine states. Baseline CBF exhibited a significant decrease after administration of caffeine (*** $p<0.001$) in both the caudal and superior GM slice, while OEF exhibited a significant increase after administration of caffeine (* $p<0.05$) in the two regions. $R_2'$ exhibited a significant increase after caffeine (* $p<0.05$) in the superior GM; no significant change was detected in baseline $CMRO_2$.

**S2. Baseline and activation changes in motor and visual ROI's**

As with the GM data (S1), the data for the unattended visual stimulus in an activation ROI were less robust than the motor data, but exhibited the same general patterns. The data for the visual ROI are shown together with the data for the motor ROI for ease of comparison. The data are summarized in Supplementary Table S2 and shown graphically in Supplementary Figure S2.

**Supplementary Table 2: Average ROI changes to caffeine**. Asterisks indicate significant difference from zero with 1-tailed T-testing. *** p<0.001; ** p<0.01; * p<0.05.

| Visual ROI | | | | | | | | |
|---|---|---|---|---|---|---|---|---|
| | **Baseline CBF** (ml/100ml/min) | **Baseline OEF** | **Baseline $R_2'$** ($sec^{-1}$) | **Baseline $CMRO_2$** (ml/100ml/min) | **δCBF to stimulus** | **δBOLD to stimulus** | **$\delta CMRO_2$ to stimulus** | λ ($\delta CMRO_2$ / δCBF) |
| Pre-caffeine | | | | | | | | |
| Mean | 73.56 | 0.33 | 4.18 | 4.77 | 0.23 | 0.013 | 0.071 | 0.28 |
| Std. dev. | 11.45 | 0.12 | 0.59 | 1.58 | 0.082 | 0.0058 | 0.061 | 0.22 |
| CV (%) | 15.56 | 35.25 | 14.20 | 33.10 | 35.23 | 43.56 | 85.4 | 80.3 |
| Post-caffeine | | | | | | | | |
| Mean | 53.96 | 0.37 | 4.26 | 3.88 | 0.21 | 0.0091 | 0.10 | 0.47 |
| Std. dev. | 13.38 | 0.21 | 1.07 | 2.08 | 0.089 | 0.0061 | 0.047 | 0.17 |
| CV (%) | 24.79 | 56.25 | 25.05 | 53.70 | 41.64 | 67.12 | 46.7 | 37.0 |
| Δ (Post-pre) | | | | | | | | |
| Mean | -19.60*** | +0.045 | +0.072 | -0.89 | -0.017 | -0.0043*** | +0.030 | +0.16* |
| Std. dev. | 10.09 | 0.21 | 1.13 | 2.07 | 0.09 | 0.0025 | 0.09 | 0.27 |
| % Δ | -26.6*** | +13.6 | +1.72 | -18.7 | -7.39 | -33.1*** | +42.3 | +57.1* |

| Motor ROI | | | | | | | | |
|---|---|---|---|---|---|---|---|---|
| | **Baseline CBF** (ml/100ml/min) | **Baseline OEF** | **Baseline $R_2'$** ($sec^{-1}$) | **Baseline $CMRO_2$** (ml/100ml/min) | **δCBF to stimulus** | **δBOLD to stimulus** | **$\delta CMRO_2$ to stimulus** | λ ($\delta CMRO_2$ / δCBF) |
| Pre-caffeine | | | | | | | | |
| Mean | 45.09 | 0.37 | 3.57 | 3.29 | 0.24 | 0.011 | 0.080 | 0.32 |
| Std. dev. | 6.79 | 0.13 | 0.69 | 1.11 | 0.039 | 0.0035 | 0.047 | 0.15 |
| CV (%) | 15.07 | 34.77 | 19.40 | 33.72 | 16.34 | 30.66 | 59.6 | 47.8 |
| Post-caffeine | | | | | | | | |
| Mean | 37.22 | 0.41 | 4.29 | 3.13 | 0.21 | 0.0083 | 0.11 | 0.47 |
| Std. dev. | 6.55 | 0.18 | 0.51 | 1.63 | 0.081 | 0.0029 | 0.063 | 0.17 |
| CV (%) | 17.59 | 44.90 | 11.94 | 52.08 | 38.16 | 34.85 | 59.8 | 35.5 |
| Δ (Post-pre) | | | | | | | | |
| Mean | -7.87*** | +0.041 | +0.71** | -0.16 | -0.028 | -0.0032** | +0.026 | +0.15*** |
| Std. dev. | 5.28 | 0.16 | 0.75 | 1.52 | 0.077 | 0.0027 | 0.060 | 0.12 |
| % Δ | -17.5*** | +10.8 | +19.9** | -4.9 | -11.7 | -29.1** | +32.5 | +46.9*** |

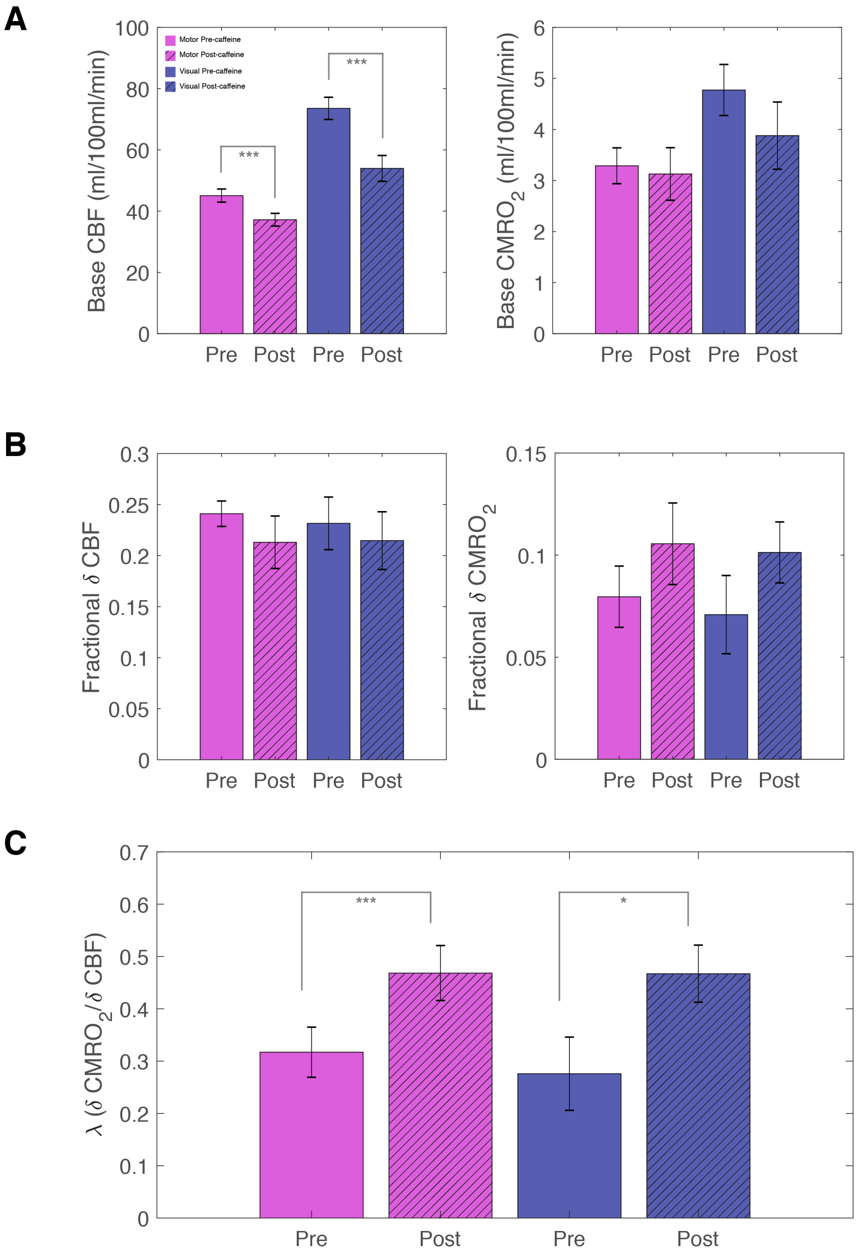

**Supplementary Figure S2: Physiological summary of changes to an intervention**. The suite of tools described in this study allow for complete quantification of physiological parameters in the baseline state, as well as changes to a stimulus. These measurements reveal the ability to discriminate between two distinct brain states, in this case before and after caffeine administration. Those physiological parameters include (**A**) baseline CBF and $CMRO_2$, (**B**) fractional $\delta$CBF and $\delta CMRO_2$ to a stimulus, and (**C**) the coupling of $\delta CMRO_2/\delta$CBF, or $\lambda$. While this study showed a significant change in baseline CBF and coupling ratio $\lambda$ from one state to the other, future applications of these methods may demonstrate other distinguishing differences. *** $p<0.001$, * $p<0.05$. Colors and cross-hatching legend in panel (**A**) indicate ROI and state.

**S3. Separation of pre-caffeine and post-caffeine states on an individual basis**

As a preliminary test for future clinical applications, we tested the extent to which these quantitative physiological methods could distinguish between the pre-caffeine and post-caffeine state in individual subjects. The goal was not to develop a test for caffeine, but rather to use caffeine as a test case for the broader question of whether these methods can distinguish a moderate change in a subject's brain state. For the caffeine challenge, the three parameters that showed the change with the highest statistical significance were scaled $\delta$BOLD/$R_2'$, baseline CBF, and the ratio of the fractional $CMRO_2$ and CBF changes to the stimulus ($\lambda$), a biomarker of neurovascular coupling. All the individual subject data were plotted as scatterplots comparing two variables (Figure S3A and C), with ellipses indicating the 95% confidence interval drawn around the mean. The parameters that were able to distinguish the two states were determined by the presence of a 95% confidence interval that did not pass through the x and y axes, which would indicate $\Delta$=0.

Specificity of these three parameters (scaled $\delta$BOLD/$R_2'$, $\lambda$, and baseline CBF) in detecting a change in brain state from pre- to post-caffeine was further tested by examining these relationships in a second set of pre-caffeine data acquisitions. While FLAIR-GESSE and VSEAN were only acquired once pre-caffeine, the functional stimulus acquisition was performed twice pre-caffeine. The first pre-caffeine run was used in comparing between pre- and post-caffeine, as its acquisition circumstances were most similar to those post-caffeine (timing after entrance into the scanner, no habituation likely). Pre- to pre-caffeine differences are shown in Figure S3B and D. While no significant change in $\lambda$ or baseline CBF was seen, as would be expected, surprisingly $\delta$BOLD/$R_2'$ was significantly different between the two pre-caffeine states. In short, the BOLD responses with repeated measurements may vary due to other physiological factors or the state of the subject, while the physiological relationships between CBF and $CMRO_2$ were more stable.

The spread of the individual pre- and post-caffeine data are plotted in Figure S4, with ellipses indicating 2 standard deviations drawn around the mean. These ellipses would surround 95% of the population. The significant overlap in range of the two populations—pre- and post-caffeine—show that a single measurement in an unknown state is not enough to identify the state itself; instead, taking two separate measurements and quantifying the difference is more demonstrative of whether a change in state has occurred.

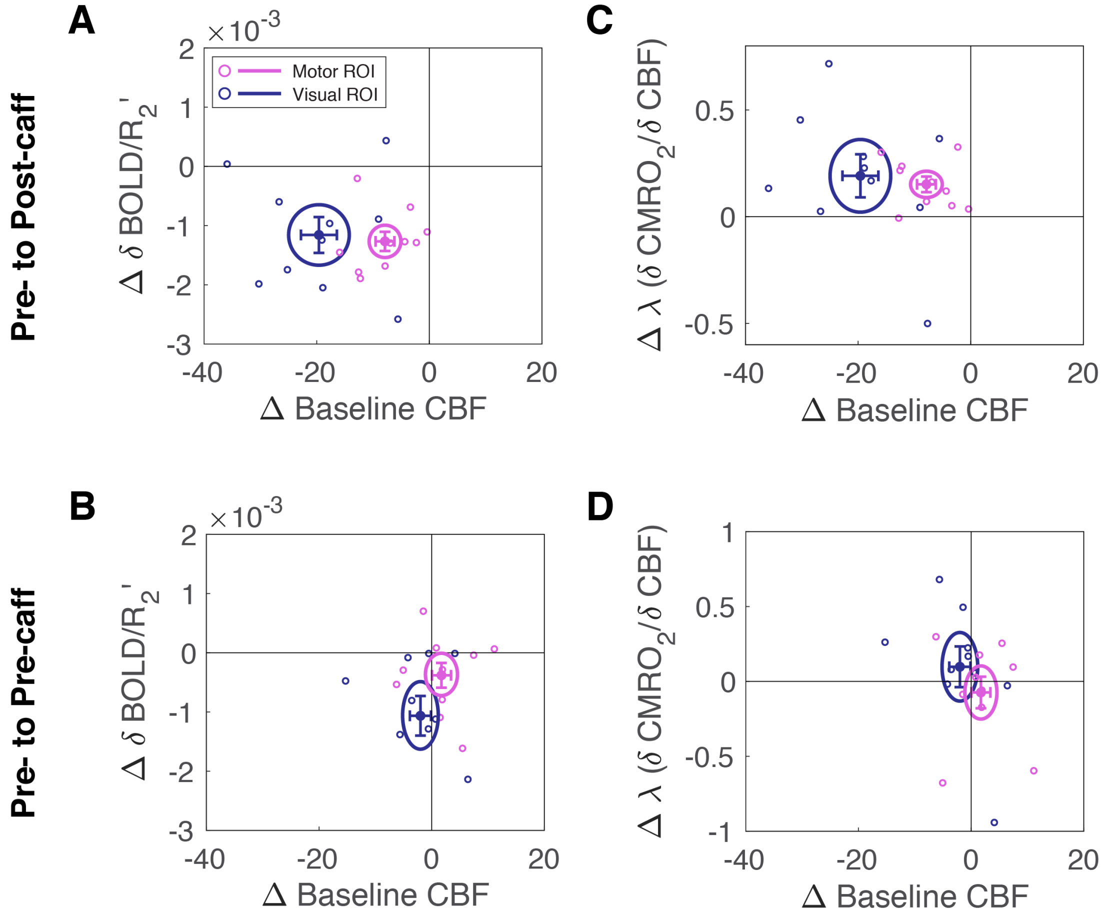

**Supplementary Figure S3: Separation of pre- and post-caffeine states**. (**A**) and (**C**) depict the major parameters distinguishing the two different conditions, pre- and post-caffeine. (**B**) and (**D**) depict the corresponding comparisons between two pre-caffeine experiments. The results in (**B**) demonstrate that δBOLD is not discriminating in its ability to indicate two different states; rather, λ (δ$CMRO_2$/δCBF) is more specific to separating the two conditions. Ellipses indicate 95% confidence intervals, error bars indicate standard error.

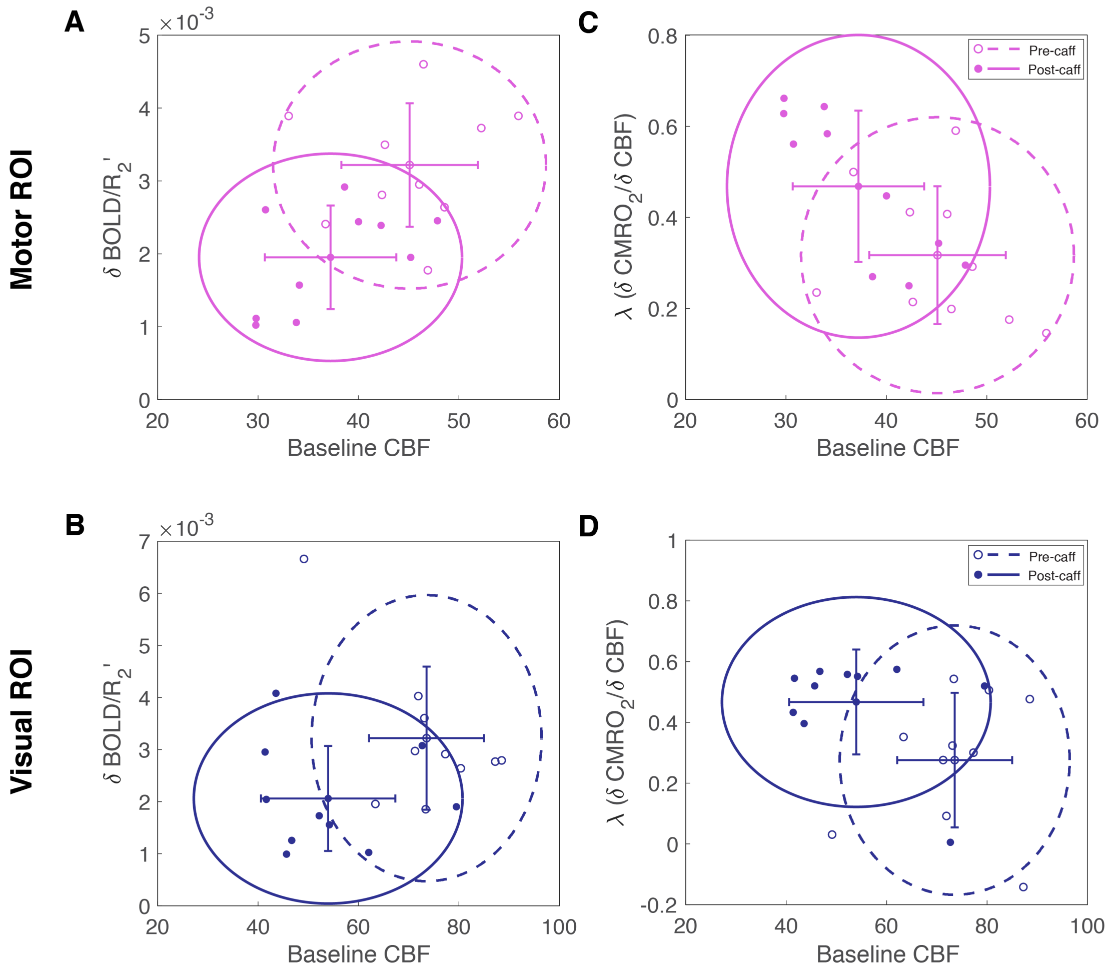

**Supplementary Figure S4: Individual data in pre- and post-caffeine states**. The scatter of individual measurements for the plotted parameters, with ellipses indicating 2 standard deviations from the mean (95% of the population), and error bars indicating standard deviation. For both the motor ROI (**A** and **C**) and visual ROI (**B** and **D**), there is significant overlap between the two states, demonstrating intersubject variability and thus the inability of any one set of measurements to designate the correct state.
